# High-yield production of recombinant human myelin oligodendrocyte glycoprotein in SHuffle bacteria without a refolding step

**DOI:** 10.1101/2024.07.22.602974

**Authors:** Wesley Wu, Sasha Gupta, Sharon A. Sagan, Carson E. Moseley, Scott S. Zamvil, John E. Pak

**Affiliations:** Chan Zuckerberg Biohub – San Francisco, San Francisco, CA, 94158, USA; Department of Neurology, Weill Institute for Neurosciences, University of California, San Francisco, San Francisco, CA, 94158, USA; Program in Immunology, University of California, San Francisco, 94143, USA

**Keywords:** Myelin oligodendrocyte glycoprotein (MOG), experimental autoimmune encephalomyelitis (EAE), autoantigen, purification, SHuffle cells, B cells

## Abstract

Experimental autoimmune encephalomyelitis (EAE) is a model for central nervous system (CNS) autoimmune demyelinating diseases such as multiple sclerosis (MS) and MOG antibody-associated disease (MOGAD). Immunization with the extracellular domain of recombinant human MOG (rhMOG), which contains pathogenic antibody and T cell epitopes, induces B cell-dependent EAE for studies in mice. However, these studies have been hampered by rhMOG availability due to its insolubility when overexpressed in bacterial cells, and the requirement for inefficient denaturation and refolding. Here, we describe a new protocol for the high-yield production of soluble rhMOG in SHuffle cells, a commercially available *E. coli* strain engineered to facilitate disulfide bond formation in the cytoplasm. SHuffle cells can produce a soluble fraction of rhMOG yielding >100 mg/L. Analytical size exclusion chromatography multi-angle light scattering (SEC-MALS) and differential scanning fluorimetry of purified rhMOG reveals a homogeneous monomer with a high melting temperature, indicative of a well-folded protein. An *in vitro* proliferation assay establishes that purified rhMOG can be processed and recognized by T cells expressing a T cell receptor (TCR) specific for the immunodominant MOG_35-55_ peptide epitope. Lastly, immunization of wild-type, but not B cell deficient, mice with rhMOG resulted in robust induction of EAE, indicating a B cell-dependent induction. Our SHuffle cell method greatly simplifies rhMOG production by combining the high yield and speed of bacterial cell expression with enhanced disulfide bond formation and folding, which will enable further investigation of B cell-dependent EAE and expand human research of MOG in CNS demyelinating diseases.

## 1.2 INTRODUCTION

The experimental autoimmune encephalomyelitis (EAE) animal model for central nervous system (CNS) demyelinating diseases, including multiple sclerosis (MS) and MOG antibody associated disease (MOGAD), is a powerful tool for studying disease pathogenesis and potential therapeutics^1,2^. Immunization with myelin oligodendrocyte glycoprotein (MOG) peptides or the recombinant extracellular domain of MOG (known as recombinant human MOG, rhMOG) is a common method for induction of EAE in mice and other animals^3,4^. The response to immunization with MOG peptides is driven by T cells and is B cell-independent^3^, referred to as T-dependent MOG EAE^2^. In contrast, the response to immunization with rhMOG requires protein processing by B cells^5,6^, and is referred to as T-B-dependent MOG EAE^2^. B cell-dependent models for MS are increasingly important given the success of monoclonal antibody-directed B cell depletion therapies in treating MS^7,8^. rhMOG is therefore a crucial resource for MS research and efficient methods for its production are needed.

Multiple approaches have been developed to produce MOG protein for T-B-dependent MOG EAE models. Initially, endogenous MOG protein was purified directly from human CNS white matter^9^, but at low yields owing to the small amount of MOG in the CNS. Later, soluble MOG was produced using one of two general approaches: a moderate yield baculovirus-mediated insect cell expression system^4^ or a high yield bacterial cell expression system that requires denaturation and refolding to produce soluble MOG.^9–12^

The MOG ectodomain adopts an IgV-like fold comprised of a sandwich of two antiparallel β-sheets^11,12^. A single, buried disulfide bond that connects the two β-sheets has been shown to significantly stabilize the IgV-like fold^13^ and, in MOG, the redox state of the disulfide bond is important for defining antigen presentation and thus whether EAE requires B cells^14^. With this in mind, we hypothesized that *E. coli* could produce large quantities of soluble, properly folded rhMOG for EAE animal studies, without requiring denaturation and refolding, if the stabilizing disulfide bond of rhMOG could be formed during protein folding inside the bacterial cell. In this study, we expressed, purified, and characterized soluble rhMOG produced in SHuffle cells, a commercially available *E. coli* strain engineered to enable disulfide bond formation in the cytoplasm^15^, achieving very high titers of EAE-inducible protein, in an efficient manner without requiring denaturation or refolding.

## 1.3 MATERIALS AND METHODS

### 1.3.1 Construct Design and Protein Expression in SHuffle T7 E.coli cells

The DNA sequence for 1-120 rhMOG (residues 30-149 from NP_996532.2) modified to include an N-terminal start codon, a C-terminal 6x His tag, and a stop codon, was optimized for *E. coli* expression using GenSMART™ Codon Optimization (https://www.genscript.com/gensmart-free-gene-codon-optimization.html). The optimized sequence was synthesized and cloned into pET-29b(+), using the NdeI and XhoI sites, by Twist Biosciences (South San Francisco, California, USA). The synthesized plasmid was transformed into SHuffle® T7 Express lysY Competent E. coli (New England Biolabs, Ipswich, Massachusetts, USA) following the manufacturer’s guidelines. For optimal soluble rhMOG expression at 1 L scale, a single transformed colony was first used to inoculate 20 mL of LB supplemented with 50 μg/mL kanamycin and grown overnight at 30 °C and 250 rpm. The 20 mL starter culture was used to inoculate 1 L of Super Broth media supplemented with 50 μg/mL kanamycin in a 2.5 L Ultra-Yield flask (Thomson Instrument Company, Oceanside, California, USA) and was grown at 37 °C and 250 rpm. The 1 L culture were grown to a high density (OD_600_= 2.0), induced with 1 mM IPTG, and shifted to 25 °C for 20 hr. Cells were harvested by centrifugation (5000g, 30 min), yielding ∼10 g of wet cell pellet, and frozen for future purification.

### 1.3.2 Purification of rhMOG

The frozen pellet was thawed on ice and resuspended in 80 mL of Buffer A (20 mM Sodium phosphate, 300 mM NaCl, pH 7.4) supplemented with 1 mg/mL lysozyme, 10 mM imidazole, 1 uL benzonase nuclease, 1 mM MgCl_2_, and 2 cOmplete EDTA-free protease inhibitor tablets (Sigma-Aldrich), followed by incubation for 30 min at 4°C with rotation. The resuspended cell mixture was then placed in an ice water bath and lysed using a Sonic Dismembrator (Fisherbrand™ Model 50, FB50110) for 3 minutes (6 cycles of 30 sec on and 30 sec off). The cell debris was pelleted by centrifugation (8000 g, 30 min) and the supernatant was loaded on a 5 mL HisTrap FF affinity chromatography column (Cytiva, Marlborough, Massachusetts, USA) using an AKTA Pure (Cytiva). The column was washed with 50 column volumes (CV) of buffer A with 20 mM imidazole and rhMOG was eluted with 20 CVs of Buffer A with 250 mM imidazole. rhMOG-containing fractions were pooled and dialyzed into either PBS or 25 mM acetate buffer, pH 4.1 using 2K or 3.5K MWCO Slidealyzer G3 dialysis cassettes (Thermo Fisher Scientific, Waltham, Massachusetts, USA). Similar results were obtained using gravity flow chromatography with 2-8 ml of 50% HisPur NiNTA resin (Thermo Fisher Scientific) in place of the AKTA Pure.

### 1.3.3 SDS-PAGE and western blot analysis of purified rhMOG

rhMOG was electrophoretically separated under reducing conditions using Mini-Protean 4-15% TGX Precast gels and a Mini-Protean Tetra Cell (Biorad, Hercules, California, USA). For total protein staining, gels were incubated with InstantBlue Coomassie Stain (Abcam) for 60 min and washed with water for 1 min. For western blot analyses, gels were transferred to PVDF membranes and blocked for 60 min with PBS + 1% BSA. The primary antibody, NYRMOG mouse monoclonal antibody (item # sc-73330, Santa Cruz Biotechnology, Dallas, Texas, USA), was used at 1:3000 dilution in Tris-buffered Saline and 0.5% Tween 20 for 1 hr. The secondary antibody, an alkaline phosphatase conjugated goat anti-mouse secondary antibody (item # 31320, Thermo Fisher Scientific), was used at 1:10,000 dilution for 1 hr. Blots were incubated with Western Blue Stabilized Substrate for Alkaline Phosphatase (Promega) for 2-5 min.

### 1.3.4 SEC-MALS of purified rhMOG

The SEC-MALS instrumentation used in this study has been described^16^. 10 μL of purified rhMOG at 1mg/mL was injected onto a Superdex 200 Increase 3.2/300 column (Cytiva, column volume = 2.4 mL) equilibrated in PBS at 0.15 mL/min. Molecular weight was determined using Astra software and graphed using Graphpad Prism. In place of using the refractive index (RI) rhMOG peak to determine protein mass (as is typical for SEC-MALS), we used the UV280 rhMOG peak due to overlap of the rhMOG RI peak with the included volume solute RI peak.

### 1.3.5 Differential scanning fluorescence of rhMOG

Differential scanning fluorescence was performed using a Prometheus NT-48 (NanoTemper). Purified rhMOG at 1 mg/mL, pre-incubated in the presence or absence of 10 mM DTT for 5 min at room temperature, was loaded into Prometheus Standard Capillaries (NanoTemper, PR-C002). Intrinsic tryptophan fluorescence was monitored at 90% excitation power. Thermal melting of rhMOG was performed from 25 °C to 95 °C, starting with a 1 min equilibration step at 25 °C followed by increasing the temperature at a rate of +1 °C per minute. Melting temperature was calculated with Prometheus software as the inflection point of the ratio of fluorescence at 350 nm to 330 nm (F350/F330) versus temperature. Data was graphed as a smoothed curve (2^nd^ order smoothing, 4 neighbors) using Graphpad Prism.

### 1.3.6 In vitro cell proliferation

CD4 positive T cells were isolated by negative magnetic sorting from the splenocytes of a MOG-TCR transgenic mouse (2D2), which recognizes the p35-55 epitope of MOG. The cells were plated at 10,000 per well in a flat bottom 96 well plate in the presence of 500,000 wild type splenocytes that had been irradiated at 3000 RAD to eliminate non-antigen targeted proliferation. The cells were cultured with various concentrations of rhMOG or MOG p35-55 for 72 hours (37°C, 5% CO2) and received 1 microcurie per well of tritiated thymidine in the final 18 hours. Proliferation was assessed by [^3^H]methylthymidine incorporation into harvested cells as previously described^17^ and is expressed as stimulation index (SI) (cpm of MOG wells/cpm of medium-only wells).

### 1.3.7 Mice

Wild type (WT) female C57BL/6 (B6), and muMT female mice (6–8 weeks of age) were purchased from Jackson Laboratory. Mice were housed in a 1:1 light:dark animal facility at the Sandler Neurosciences Center of University of California, San Francisco (UCSF).

### 1.3.8 Standard Protocol Approvals and Registrations

All animal experiments were conducted in accordance with UCSF’s Institutional Animal Care and Use Committee. Standard protocol approvals and registrations were obtained (AN188984-02B).

### 1.3.9 EAE induction

Five WT and five B cell deficient mice were immunized subcutaneously with an emulsion of 100 µg of rhMOG in complete Freund’s adjuvant. 300 ng of Pertussis toxin is injected intraperitoneally at the time of the immunization and two days later. Clinical EAE symptoms are scored daily for 21 days using a clinical scoring system that describes neurological symptoms on a 0-5 scale, where 0 = no clinical disease, 1 = loss of tail tone only, 2 = mild monoparesis or paraparesis, 3 = severe paraparesis, 4 = paraplegia and/or quadriparesis, and 5 = moribund or death.

## 1.4 RESULTS

### 1.4.1 Soluble rhMOG is produced in SHuffle *E.coli* cells

The reductive environment of the *E.coli* cytoplasm is normally incompatible with disulfide bond formation^18^, and recombinant proteins that have disulfide bonds in their native conformation often form insoluble inclusion bodies when overexpressed in *E.coli*^19^. The oxidative environment and disulfide bond formation (Dsb^20^) proteins of the periplasm can promote disulfide bonding, however, achieving high titers of recombinant protein in the periplasm is non-trivial^21^. SHuffle *E.coli* cells^15^, with their ability to promote disulfide bond formation in the cytoplasm, could potentially overexpress high titers of soluble, well-behaved rhMOG given that the buried disulfide bond of rhMOG (Figure 1A, B) may contribute to its stability and/or folding.

**Figure 1.**
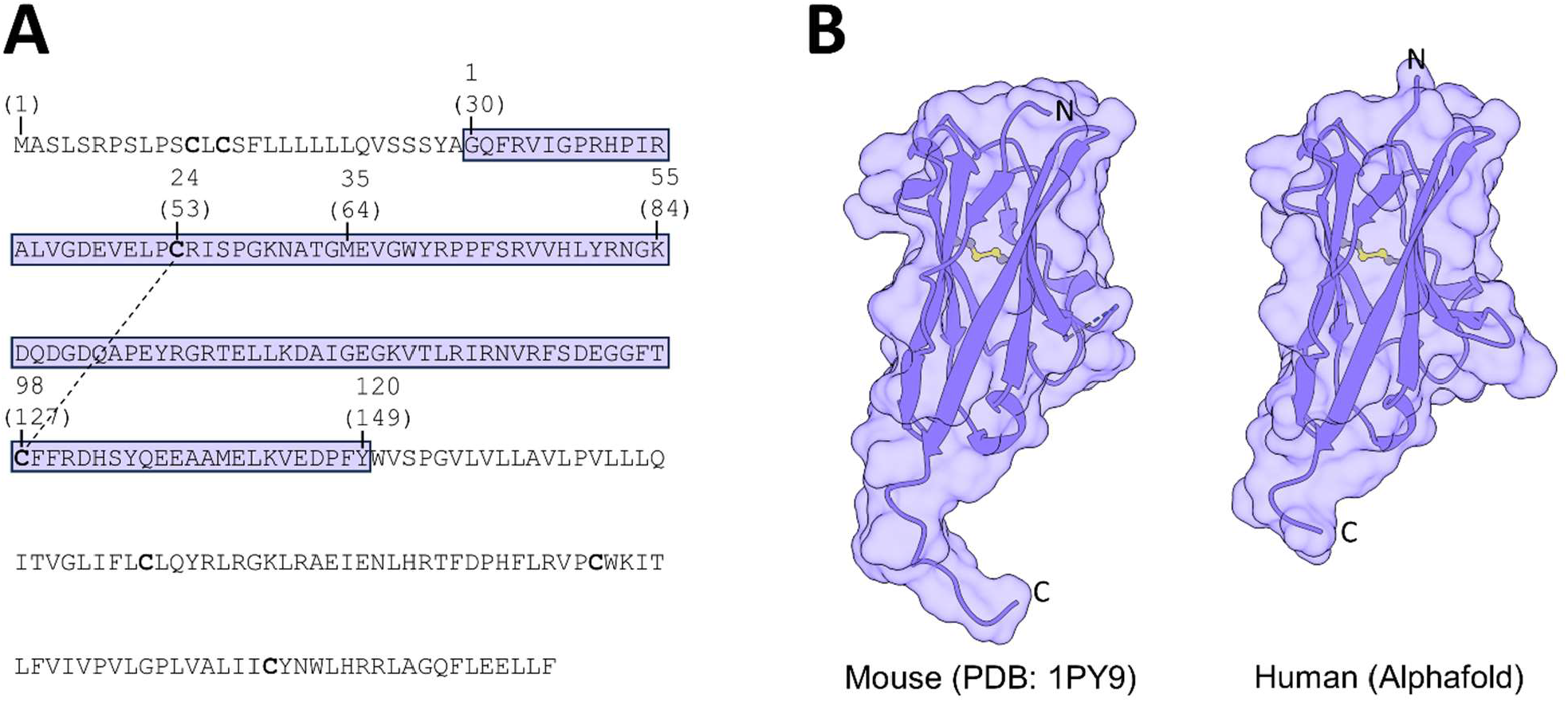
Sequence and structure of MOG. A) Amino acid sequence of human MOG. rhMOG is highlighted in purple with residues numbered in the absence and presence (in parentheses) of signal peptide. The disulfide bond of rhMOG is denoted with a dashed line. All cysteine residues of MOG are denoted in bold. B) Comparison of mouse MOG and rhMOG structures. Disulfide bonds are shown in ball-and-stick representation (sulfur atom in yellow). The rhMOG structure was predicted using Alphafold2^22^. Structure renderings were created using ChimeraX^23^.

To evaluate if promoting disulfide bond formation could improve rhMOG production in bacteria, we first performed small-scale expression tests in SHuffle *E. coli* and BL21(DE3) *E.coli* cells grown in Luria Broth (LB). The sequence of human rhMOG (Figure 1A, Supplemental Figure 1), modified to include an N-terminal methionine and a C-terminal His_6_ tag, was codon optimized, synthesized, and cloned into a pET-29a T7 bacterial expression vector. After inducing expression for 4 hr at 37 °C, soluble and insoluble bacterial fractions were analyzed by SDS-PAGE (Figure 2A). Both bacterial strains showed robust expression of rhMOG (expected MW=14.7 kDa), however, only SHuffle extracts showed a significant amount of rhMOG in the soluble fraction (Figure 2A), estimated to be ∼10% of total soluble protein.

**Figure 2.**
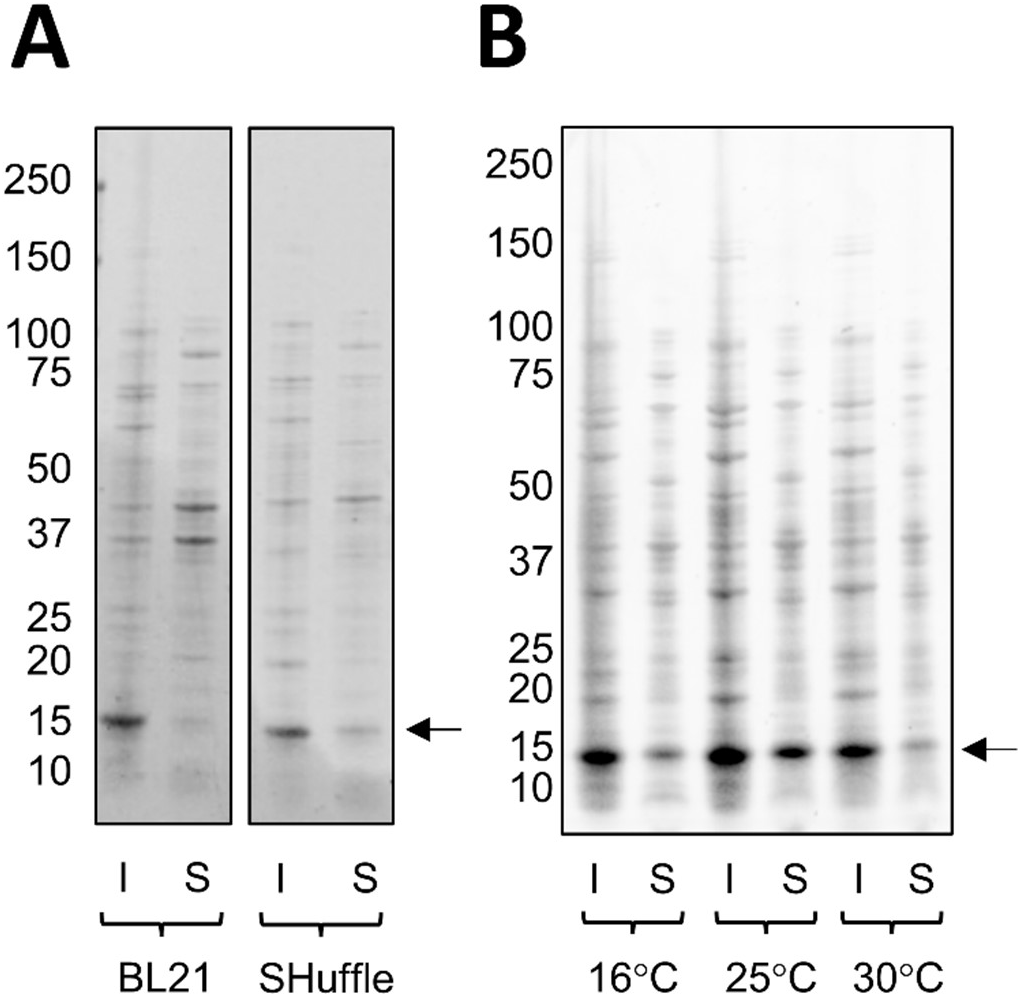
Effects of host strain and induction temperature on the expression of soluble fraction rhMOG. A) Comparison of insoluble (“I”) and soluble (“S”) lysate fractions from A) BL21 (DE3) *E.coli* rhMOG and SHuffle *E.coli* rhMOG induced at 37 °C for 4 hr and B) SHuffle *E.coli* rhMOG induced at lower temperatures for 24 hr. All samples were analyzed by reducing SDS-PAGE and stained with Coomassie blue. Arrows denote the expected migration position of rhMOG.

Encouraged by these initial results with SHuffle cells, we evaluated the effect of induction temperature on soluble rhMOG production, with an expectation that rhMOG expression titers could increase at temperatures below 37 °C^24^. Following IPTG induction of small-scale cultures, we lowered expression temperatures to 30 °C, 25 °C and 16 °C, and grew the cultures for 24 hrs to compensate for the lower rate of protein production in *E.coli* at temperatures below 37 °C^25^. While each lower temperature results in high levels of rhMOG expression (Figure 2B), cells grown at 25 °C yield a large increase in soluble rhMOG titers, estimated to be ∼30% of total soluble protein (Figure 2B).

### 1.4.2 Optimization of scaled up rhMOG expression and purification

To determine the best conditions for producing large quantities of soluble rhMOG, we evaluated several growth conditions at liter scale. Soluble rhMOG was extracted from cells by ultrasonic lysis, purified by Ni-NTA affinity purification, and stored, after dialysis, in PBS or acetate buffer. The yield of purified soluble rhMOG expressed in LB was 14 mg per L of cell culture. Substituting LB with Super Broth showed a large improvement in soluble rhMOG titers at liter scale. Purified yields of soluble rhMOG from SHuffle *E.coli* grown in Super Broth were ∼40 mg/L, 2.5-fold higher than that of LB. Furthermore, delaying IPTG induction to when the Super Broth cell cultures were at a higher density (OD_600_= 2.0) results in a further 2.5-fold increase in rhMOG titers, yielding an optimized yield of ∼100 mg of purified soluble rhMOG per L of cell culture. The purity of the final formulated rhMOG preparation is estimated to be >99% based on reducing SDS-PAGE analysis (Figure 3A).

**Figure 3.**
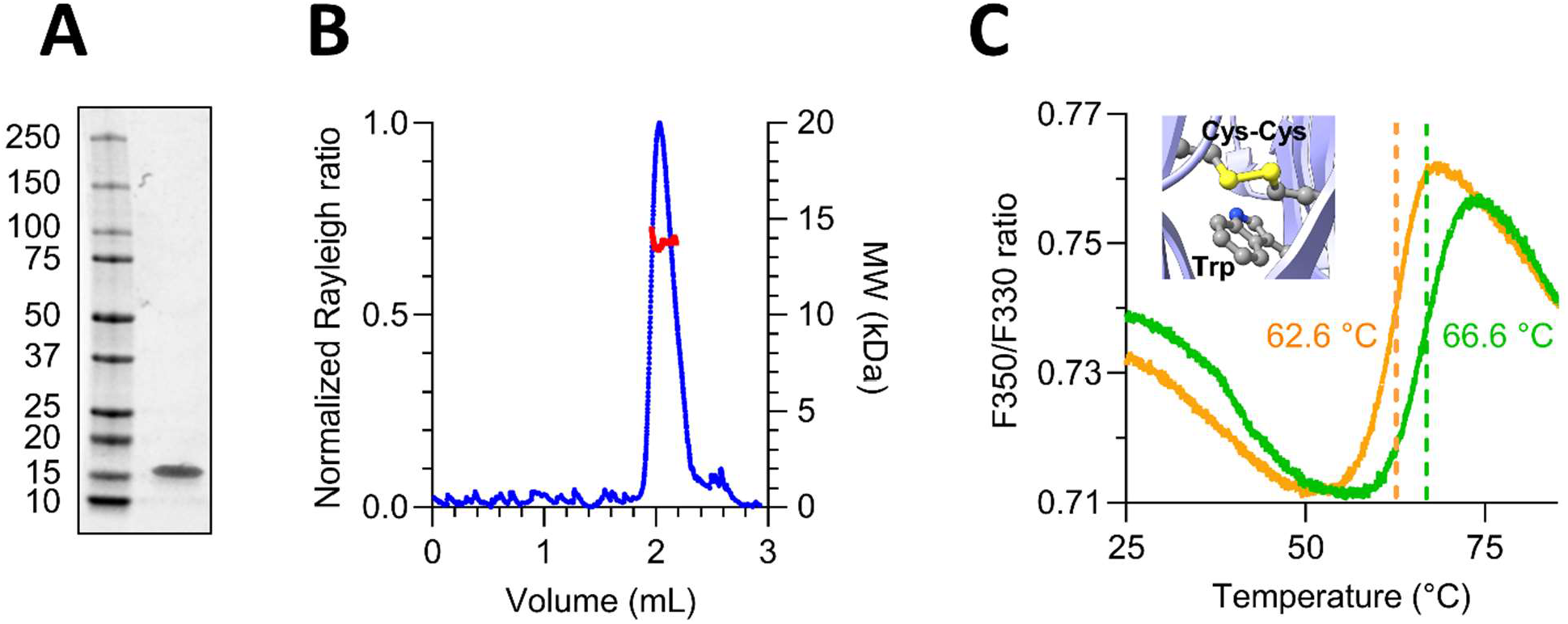
rhMOG is a pure, homogeneous and thermostable protein. A) Reducing SDS-PAGE of rhMOG stained with Coomassie Blue. B) SEC-MALS analysis of rhMOG. Sample was separated on a Superdex 200 Increase 3.2/300 column at 0.15 mL/min in PBS. Normalized light scattering (blue, left axis) and calculated molecular weight (red, right axis) are shown. C) Effect of reducing agent on the thermostability of rhMOG. Differential scanning intrinsic tryptophan fluorescence of rhMOG in the absence (green) and presence (orange) of 10 mM DTT. Melting temperature (T_m_) is denoted with a dashed line. Insert shows a close-up view of the structure of human MOG (Alphafold2 model) in the same orientation as that shown in Figure 1B). The sole tryptophan and disulfide bond of rhMOG are shown in ball-and-stick representation (grey carbon, blue nitrogen, yellow sulfur).

### 1.4.3 rhMOG purified from SHuffle *E. coli* is monomeric and thermostable

To confirm that the soluble rhMOG purified from SHuffle *E.coli* was well folded and monodisperse, we measured its oligomeric state by size-exclusion chromatography with multi-angle light scattering (SEC-MALS). A single light scattering peak is resolved on a Superdex 200 Increase 3.2/300 size exclusion column, corresponding to a monomer with a calculated molecular weight of ∼14 kDa (Figure 3B), consistent with that reported for recombinant, refolded MOG ectodomain in solution^11,12^. Light scattering does not reveal oligomerization or aggregates that would be a hallmark for incorrect intermolecular disulfide bonding or poorly folded rhMOG.

The well-behaved SEC-MALS peak does not necessarily indicate that the functionally important^14^ and putatively stabilizing^13^ disulfide bond of rhMOG has formed, given that the IgV-like fold adopts a similar overall conformation with or without its disulfide bond^13^. To quantify the effect of disulfide bonding on rhMOG stability, we determined the thermal melting temperature (T_m_) of rhMOG by differential scanning fluorimetry (DSF), in the absence and presence of reducing agent (Figure 3C). The sole tryptophan of rhMOG is buried (Figure 3C) and is ideally situated to report on unfolding. The purified rhMOG preparation, which does not include reducing agent in the final formulation, undergoes a cooperative transition with a high T_m_ of 66.6 °C (Figure 3C). In the presence of 10mM DTT, which would reduce any disulfide bond that is present, the protein is comparatively destabilized with a lower T_m_ of 62.6 °C (Figure 3C). Taken together, we show that soluble rhMOG that is purified from SHuffle *E.coli*, without requiring denaturation and refolding, does indeed contain a stabilizing disulfide bond that likely enables high levels of soluble rhMOG expression.

### 1.4.4 rhMOG purified from SHuffle *E.coli* stimulates proliferation of MOG-specific T cells *in vitro*

EAE induced by rhMOG protein, in contrast to MOG 35-55 peptide, requires B cells^5,6^. The processing and presentation of MOG epitopes to T cells, nevertheless, is characteristic of both T-dependent and T-B-dependent MOG EAE^5^. To determine if rhMOG purified from SHuffle *E.coli* could be processed to present MOG epitopes to T cells, we measured MOG-specific T cell proliferation *in vitro*^26^. CD4+ T cells were isolated from MOG 35-55-specific TCR (2D2) transgenic (Tg) mice^26^ and cultured with irradiated splenocytes. In the presence of MOG 35-55 peptide or purified rhMOG protein, which was confirmed by western blotting (Figure 4A), T cell proliferation is stimulated in a dose dependent manner (Figure 4B). Thus, rhMOG purified from SHuffle *E.coli* can indeed be efficiently taken up and processed by APCs, resulting in robust stimulation of CD4+ T cells that are specific for the MOG 35-55 epitope.

**Figure 4.**
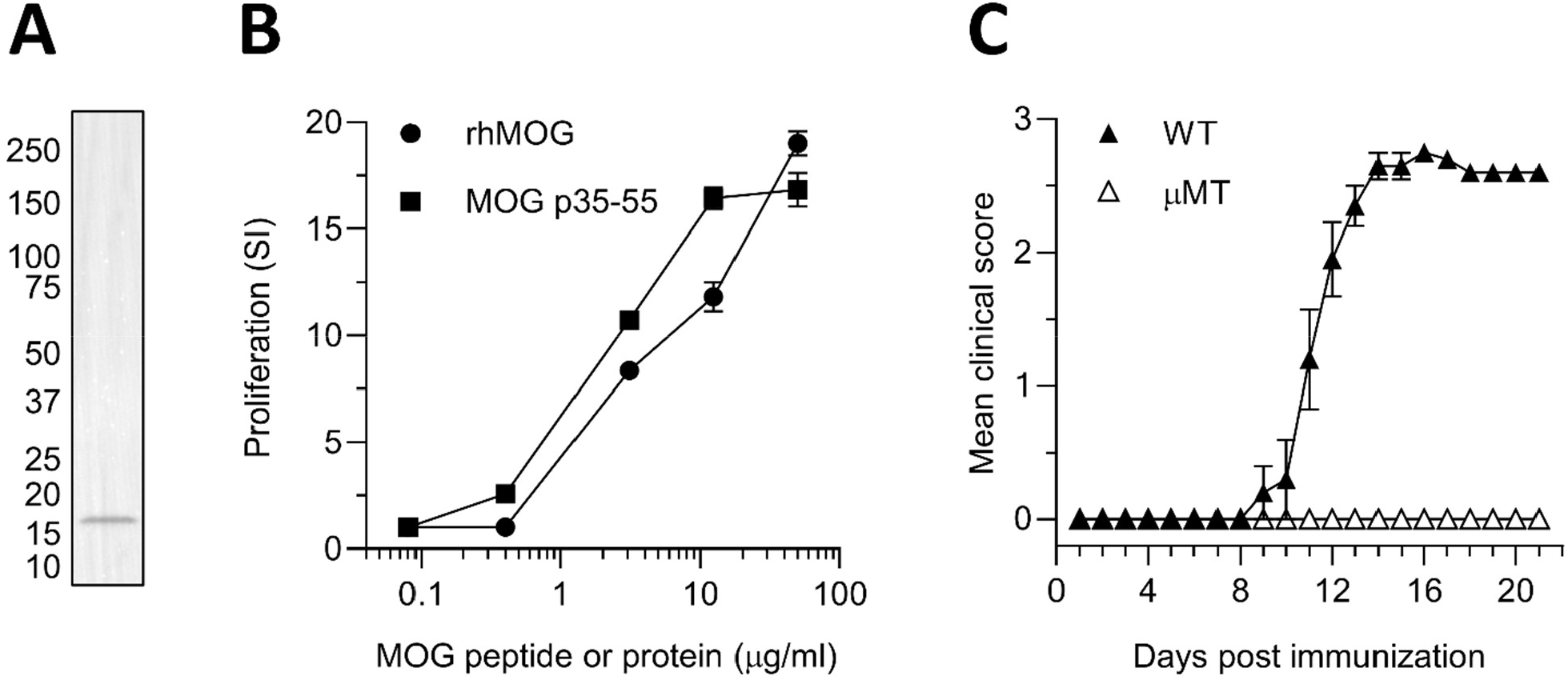
Functional activity of rhMOG *in vitro* and *in vivo*. A) Western blot of purified rhMOG using MOG 35-55-specific primary antibody NYRMOG. B) *In vitro* proliferation of CD4+ T cells isolated from 2D2 transgenic mice and stimulated with MOG 35-55 peptide (filled square) or rhMOG protein (filled circle). Mean proliferation, expressed as stimulation index (SI), and standard error of the mean (SEM) are shown. C) *In vivo* induction of B cell-dependent EAE in mice. Clinical score was determined daily for wild type (“WT”, filled triangle) and B cell deficient (“μMT”, open triangle) mice that were immunized with rhMOG. Mean clinical score and SEM are shown.

### 1.4.5 rhMOG purified from SHuffle *E.coli* induces B cell-dependent EAE in mice

While rhMOG purified from SHuffle *E.coli* is a soluble and well-folded protein that induces T cell proliferation *in vitro*, it was important to explore the *in vivo* effect, and to determine if our rhMOG could induce EAE and do so in a B cell-dependent manner. This is particularly important as poor stability of rhMOG, *in vivo*, could lead to proteolytic degradation of rhMOG that would circumvent the need for B cells in the processing and presentation of MOG epitopes.

To determine if our rhMOG could induce EAE in a B cell-dependent manner, we immunized wild type (WT) and genetically B cell deficient (μMT^27^) mice with 100 μg of rhMOG protein. WT mice immunized with rhMOG showed an onset of EAE symptoms 9 days post immunization, with a maximum mean clinical score at 13-21 days post immunization, matching closely with previously reported EAE mouse models^6^. In contrast, µMT mice immunized with rhMOG showed no clinical symptoms up to 21 days post immunization, indicating that the EAE response in WT mice was B cell-dependent. Thus, our method for expressing and purifying rhMOG does indeed produce folded protein that is functional *in vivo* in a B cell-dependent manner, without the need for denaturation and refolding.

## 1.5 DISCUSSION

B cell involvement in demyelinating diseases including multiple sclerosis is evident by the tremendous success of B cell depleting therapy^7,8^, yet the most commonly used model of CNS demyelination, T-dependent MOG EAE, is B cell-independent. rhMOG is an important reagent used to generate T-B-dependent MOG EAE that until now has been restricted due to large scale production issues. In addition, properly folded rhMOG allows for the examination of autoantibodies against conformation-dependent epitopes, in contrast to MOG peptide-specific autoantibodies which do not recognize the native MOG protein^12^. Previous methods for recombinant production have required time and labor intensive baculovirus/insect cell expression protocols or refolding protocols after bacterial expression as insoluble inclusion bodies^4,9–12^. We show that expressing rhMOG in SHuffle *E.coli*, which promotes disulfide bond formation in the bacterial cytosol, enables soluble expression of rhMOG without the need for refolding from inclusion bodies. Upon scale up and optimization of induction and expression conditions, yields of over 100 mg/L of purified rhMOG were achieved through use of SHuffle *E.coli*. The rhMOG purified from SHuffle cells stimulates MOG-specific T cells in an *in vitro* proliferation assay and can be used to induce T-B-dependent MOG EAE in mice.

Recombinant expression in SHuffle *E.coli* is a highly efficient method for producing rhMOG compared to established techniques. Insect cell cultures require generating recombinant baculovirus, subsequent infection/expression, and purification of secreted protein from the media, which together can take several weeks. Insoluble bacterial cell produced inclusion bodies can attain high yields of rhMOG but require refolding steps that can add complexity and stochasticity to the purification process. Our work combines the high yields and speed of bacterial expression with the improved folding of rhMOG that is seen in insect cells, resulting in an accessible method for producing the quantities of rhMOG that are needed for T-B-dependent MOG EAE models.

## 1.6 CRedIT AUTHORSHIP CONTRIBUTION STATEMENT

**Wesley Wu:** Conceptualization, Methodology, Validation, Formal analysis, Investigation, Writing – Original Draft, Writing – Review & Editing, Visualization, Project administration. **Sasha Gupta:** Conceptualization, Validation, Writing – Review & Editing, Project administration. **Sharon A. Sagan:** Methodology, Validation, Formal Analysis, Investigation, Writing – Review & Editing, Visualization. **Carson E. Moseley:** Writing – Review & Editing. **Scott S. Zamvil**: Resources, Writing – Review & Editing, Supervision, Funding acquisition. **John E. Pak:** Formal analysis, Investigation, Writing – Original Draft, Writing – Review & Editing, Visualization, Supervision, Funding acquisition.

## 1.7 ACKNOWLEDGEMENTS

JEP and WW thank Priscilla Chan and Mark Zuckerberg for their funding support at the Chan Zuckerberg Biohub San Francisco. SSZ is supported by grants from the NIH (1 R01 AI131624-01A1 and 1 RO1 AI170863-01A1), The Sumaira Foundation and the Weill Institute for Neurosciences. CEM is supported by a Clinician Scientist Development Award Grant FAN-2107-38301 from the National Multiple Sclerosis Society, the C.D. Spangler Foundation, and a Clinician-Scientist Award from the Weill Neurohub. We thank Sandra L. Schmid for providing feedback on the manuscript.

---

**Supplemental Figure 1.**
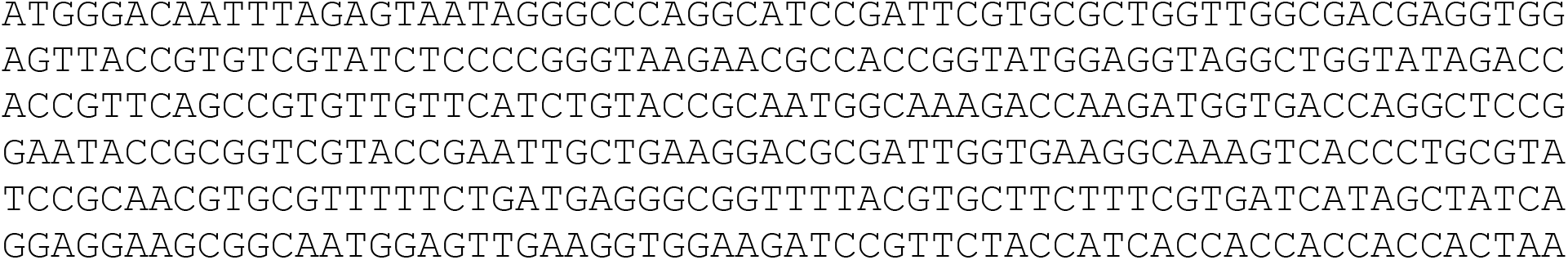
Codon-optimized nucleotide sequence of rhMOG.

## REFERENCES

(1) Constantinescu, C. S., Farooqi, N., O’Brien, K., and Gran, B. (2011) Experimental autoimmune encephalomyelitis (EAE) as a model for multiple sclerosis (MS). Br J Pharmacol 164, 1079–1106.

(2) Moseley, C. E., Virupakshaiah, A., Forsthuber, T. G., Steinman, L., Waubant, E., and Zamvil, S. S. (2024) MOG CNS Autoimmunity and MOGAD. Neurology Neuroimmunology & Neuroinflammation 11, e200275.

(3) de Rosbo, N. K., Mendel, I., and Ben-Nun, A. (1995) Chronic relapsing experimental autoimmune encephalomyelitis with a delayed onset and an atypical clinical course, induced in PL/J mice by myelin oligodendrocyte glycoprotein (MOG)-derived peptide: Preliminary analysis of MOG T cell epitopes. European Journal of Immunology 25, 985–993.

(4) Devaux, B., Enderlin, F., Wallner, B., and Smilek, D. E. (1997) Induction of EAE in mice with recombinant human MOG, and treatment of EAE with a MOG peptide. Journal of Neuroimmunology 75, 169–173.

(5) Lyons, J.-A., San, M., Happ, M. P., and Cross, A. H. (1999) B cells are critical to induction of experimental allergic encephalomyelitis by protein but not by a short encephalitogenic peptide. European Journal of Immunology 29, 3432–3439.

(6) Molnarfi, N., Schulze-Topphoff, U., Weber, M. S., Patarroyo, J. C., Prod’homme, T., Varrin-Doyer, M., Shetty, A., Linington, C., Slavin, A. J., Hidalgo, J., Jenne, D. E., Wekerle, H., Sobel, R. A., Bernard, C. C. A., Shlomchik, M. J., and Zamvil, S. S. (2013) MHC class II-dependent B cell APC function is required for induction of CNS autoimmunity independent of myelin-specific antibodies. J Exp Med 210, 2921–2937.

(7) Hauser, S. L., Waubant, E., Arnold, D. L., Vollmer, T., Antel, J., Fox, R. J., Bar-Or, A., Panzara, M., Sarkar, N., Agarwal, S., Langer-Gould, A., Smith, C. H., and HERMES Trial Group. (2008) B-cell depletion with rituximab in relapsing-remitting multiple sclerosis. N Engl J Med 358, 676–688.

(8) Hauser, S. L., Belachew, S., and Kappos, L. (2017) Ocrelizumab in Primary Progressive and Relapsing Multiple Sclerosis. N Engl J Med 376, 1694.

(9) Abo, S., Bernard, C. C., Webb, M., Johns, T. G., Alafaci, A., Ward, L. D., Simpson, R. J., and Kerlero de Rosbo, N. (1993) Preparation of highly purified human myelin oligodendrocyte glycoprotein in quantities sufficient for encephalitogenicity and immunogenicity studies. Biochem Mol Biol Int 30, 945–958.

(10) Bettadapura, J., Menon, K. K., Moritz, S., Liu, J., and Bernard, C. C. A. (1998) Expression, Purification, and Encephalitogenicity of Recombinant Human Myelin Oligodendrocyte Glycoprotein. Journal of Neurochemistry 70, 1593–1599.

(11) Clements, C. S., Reid, H. H., Beddoe, T., Tynan, F. E., Perugini, M. A., Johns, T. G., Bernard, C. C. A., and Rossjohn, J. (2003) The crystal structure of myelin oligodendrocyte glycoprotein, a key autoantigen in multiple sclerosis. Proc Natl Acad Sci U S A 100, 11059–11064.

(12) Breithaupt, C., Schubart, A., Zander, H., Skerra, A., Huber, R., Linington, C., and Jacob, U. (2003) Structural insights into the antigenicity of myelin oligodendrocyte glycoprotein. Proc Natl Acad Sci U S A 100, 9446–9451.

(13) Goto, Y., and Hamaguchi, K. (1979) The role of the intrachain disulfide bond in the conformation and stability of the constant fragment of the immunoglobulin light chain. J Biochem 86, 1433–1441.

(14) Bergman, C. M., Marta, C. B., Maric, M., Pfeiffer, S. E., Cresswell, P., and Ruddle, N. H. (2012) A switch in pathogenic mechanism in myelin oligodendrocyte glycoprotein-induced experimental autoimmune encephalomyelitis in IFN-γ-inducible lysosomal thiol reductase-free mice. J Immunol 188, 6001–6009.

(15) Lobstein, J., Emrich, C. A., Jeans, C., Faulkner, M., Riggs, P., and Berkmen, M. (2012) SHuffle, a novel Escherichia coli protein expression strain capable of correctly folding disulfide bonded proteins in its cytoplasm. Microb Cell Fact 11, 56.

(16) Puccinelli, R. R., Sama, S. S., Worthington, C. M., Puschnik, A. S., Pak, J. E., and Gómez-Sjöberg, R. (2024) Open-source milligram-scale, four channel, automated protein purification system. PLOS ONE 19, e0297879.

(17) Sagan, S. A., Cruz-Herranz, A., Spencer, C. M., Ho, P. P., Steinman, L., Green, A. J., Sobel, R. A., and Zamvil, S. S. (2017) Induction of Paralysis and Visual System Injury in Mice by T Cells Specific for Neuromyelitis Optica Autoantigen Aquaporin-4. J Vis Exp 56185.

(18) Stewart, E. J., Aslund, F., and Beckwith, J. (1998) Disulfide bond formation in the Escherichia coli cytoplasm: an in vivo role reversal for the thioredoxins. EMBO J 17, 5543–5550.

(19) Fischer, B., Sumner, I., and Goodenough, P. (1993) Isolation, renaturation, and formation of disulfide bonds of eukaryotic proteins expressed in Escherichia coli as inclusion bodies. Biotechnol Bioeng 41, 3–13.

(20) Nakamoto, H., and Bardwell, J. C. A. (2004) Catalysis of disulfide bond formation and isomerization in the Escherichia coli periplasm. Biochimica et Biophysica Acta (BBA) - Molecular Cell Research 1694, 111–119.

(21) Karyolaimos, A., and de Gier, J.-W. (2021) Strategies to Enhance Periplasmic Recombinant Protein Production Yields in Escherichia coli. Front Bioeng Biotechnol 9, 797334.

(22) Jumper, J., Evans, R., Pritzel, A., Green, T., Figurnov, M., Ronneberger, O., Tunyasuvunakool, K., Bates, R., Žídek, A., Potapenko, A., Bridgland, A., Meyer, C., Kohl, S. A. A., Ballard, A. J., Cowie, A., Romera-Paredes, B., Nikolov, S., Jain, R., Adler, J., Back, T., Petersen, S., Reiman, D., Clancy, E., Zielinski, M., Steinegger, M., Pacholska, M., Berghammer, T., Bodenstein, S., Silver, D., Vinyals, O., Senior, A. W., Kavukcuoglu, K., Kohli, P., and Hassabis, D. (2021) Highly accurate protein structure prediction with AlphaFold. Nature 596, 583–589.

(23) Pettersen, E. F., Goddard, T. D., Huang, C. C., Meng, E. C., Couch, G. S., Croll, T. I., Morris, J. H., and Ferrin, T. E. (2021) UCSF ChimeraX: Structure visualization for researchers, educators, and developers. Protein Sci 30, 70–82.

(24) Radmard, M., and Hashemi, A. (2024) Response Surface Methodology Approach to Optimize the Expression of Thioredoxin-MOG Fusion Protein. Pharm Sci 30, 252–261.

(25) Farewell, A., and Neidhardt, F. C. (1998) Effect of Temperature on In Vivo Protein Synthetic Capacity in Escherichia coli. J Bacteriol 180, 4704–4710.

(26) Bettelli, E., Pagany, M., Weiner, H. L., Linington, C., Sobel, R. A., and Kuchroo, V. K. (2003) Myelin Oligodendrocyte Glycoprotein–specific T Cell Receptor Transgenic Mice Develop Spontaneous Autoimmune Optic Neuritis. Journal of Experimental Medicine 197, 1073–1081.

(27) Kitamura, D., Roes, J., Kühn, R., and Rajewsky, K. (1991) A B cell-deficient mouse by targeted disruption of the membrane exon of the immunoglobulin mu chain gene. Nature 350, 423–426.

